# A novel function for Prdm12 during neural crest migration reveals a link between Wnt and N-cadherin

**DOI:** 10.1101/2025.07.28.667246

**Authors:** Subham Seal, Cécile Milet, Chenxi Zhou, Anne-Hélène Monsoro-Burq

## Abstract

The delamination of neural crest cells is a critical developmental event shaping the vertebrate head and peripheral nervous system, among other tissues. While the gene regulatory network driving neural crest formation (NC-GRN) has been roughly drafted, there are many fine-tuning mechanisms which require full exploration, especially when a complex cross-talk between several regulators and signaling pathways is involved. We have identified that *Prdm12*, which encodes a histone methyltransferase highly expressed in the central nervous system and lateral preplacodal ectoderm, is also expressed in the *sox10*-positive cells located at the lateral front of the premigratory neural crest domain in *Xenopus laevis* embryos. We show that Prdm12 regulates cranial neural crest emigration, independently of its known enzymatic activity, by regulating non-canonical WNT signaling, which in turn controls N-cadherin membrane localization. Our work elucidates an important function of Prdm12 in the neural crest cells initiating migration and establishes a novel epistatic link between WNT signaling pathways and cell migration in the NC-GRN.

## Introduction

The neural crest (NC) is a multipotent cell population in the vertebrate embryo, giving rise to over 30 different cell and tissue types. This amazing diversity in derivatives is accompanied by the migration of NC cells to distant sites in the embryo. Ectodermal in origin, the epithelial-like NC cells first undergo a process called epithelial-to-mesenchymal-transition (EMT), enabling them to perform cellular migration. During EMT, the NC cells lose their epithelial characteristics (apico-basally polarized, non-motile cells grouped together into a layer through the presence of tight junctions) to become more mesenchymal (front-back polarized, loose, spindle-shaped cells with invasive capabilities; Campbell and Casanova, 2016; Nieto et al., 2016). In most vertebrate species, the NC cells undergo EMT during mid-to-late neurula stages, although the start of NC migration differs (Leathers and Rogers, 2022). Interestingly, the process of EMT shares multiple parallels with cancer metastasis, underscoring its broader biological relevance (Brabletz et al., 2018; Huang et al., 2022; Kerosuo and Bronner-Fraser, 2012).

EMT is broadly regulated by a conserved cohort of specific factors (including Snai1/2, Twist1 and Zeb1/2 transcriptional regulators), collectively termed as EMT-TFs, which form a key module of the NC gene-regulatory-network (NC-GRN): EMT-TF depletion impairs NC-EMT and subsequently leads to perturbed NC migration and differentiation (Fazilaty et al., 2019; Martik and Bronner, 2017). During NC cell EMT, most of the EMT-TFs directly affect other cellular components either by gene expression regulation or protein-protein interactions. For example, while Snai2 or Zeb2 reduce the level of *cdh1* (*e-cadherin)* expression (Rogers et al., 2013; Tien et al., 2015), Twist1 directly interacts with catenins to modulate cytoskeletal dynamics (Bertol et al., 2022). Recently, studies have also focused on establishing the cooperation between these EMT-TFs themselves (Cao, 2024; Guzman-Espinoza et al., 2024; McDermott et al., 2025; Waryah et al., 2023; Youssef et al., 2024).

NC migration is further regulated by several other molecules and pathways. Structural components like cadherins (Cdh1, Cdh2, Cdh11) and metalloproteinases (MMPs and ADAMs) regulate delamination of NCCs by dissociating the epithelial layer and degrading the extracellular matrix (Abbruzzese et al., 2016; Ahsan et al., 2019; Coles et al., 2007; Dady et al., 2012; Huang et al., 2016; Kashef et al., 2009; Langhe et al., 2016; Mathavan et al., 2017; McCusker et al., 2009; Rogers et al., 2018; Scarpa et al., 2015; Schiffmacher et al., 2016). A few metalloproteinases, such as MMP2/9 and MMP14/28, also feedback to regulate the levels of EMT factors and cadherins (Garmon et al., 2018; Gouignard et al., 2023; Monsonego-Ornan et al., 2012; Schiffmacher et al., 2016). Subsequently, polarity components like Par3, Rho, Rac and others, ensure the timing and directionality of migration, by establishing a front (Rac1+) to back (RhoA+) polarity (Fort and Théveneau, 2014; Gossen et al., 2024; Grund et al., 2021; Podleschny et al., 2015). In the context of signaling pathways, the chick NC requires high BMP signaling and inhibition of FGF and Wnt signaling for NC delamination (Hutchins and Bronner, 2018; Martínez-Morales et al., 2011; Piacentino and Bronner, 2018; Zhang et al., 2018). Interestingly, the frog NC requires Wnt signaling, although maintaining the delicate balance between the canonical and non-canonical Wnt pathways is crucial (Gonzalez Malagon et al., 2018; Maj et al., 2016).

In sum, multiple studies have uncovered different molecular factors regulating NC EMT and migration. However, these studies have mostly been performed on individual genes and therefore, how these multiple players interact and cooperate during this process remains poorly understood. This has resulted in many missing connections within the NC-GRN, especially to link transcriptional to signaling activity, and to cellular mechanisms. To strengthen our current understanding of the NC-GRN (Hovland et al., 2019; Martik and Bronner, 2017), we searched our previous RNAseq datasets (Kotov et al., 2024; Plouhinec et al., 2014) for potential novel players. We identified *prdm12*, encoding a histone methyltransferase, as a target of the neural border (NB) specifier Pax3, a pivotal regulator of the NC-GRN (Monsoro-Burq et al., 2005). Interestingly, Prdm12 has previously been implicated in the regulation of NC derivatives formation: in mouse embryos, the reduction of Prdm12 levels, which is co-expressed with Sox10 before neurogenesis, depletes the Sox10+ NC cells at E12.5 (Bartesaghi et al., 2019). Although this does not mean that Prdm12 regulates Sox10 directly, it suggests that Prdm12 promotes proliferation and maintenance of the NC cell lineages. Similarly, Prdm12 is necessary for the specification of skin NC stem cells into TrkA+ nociceptors (Bataille et al., 2020). Together, this suggested a potential role of Prdm12 during NC development.

A previous study in the *Xenopus laevis* NC has shown that Prdm12, which is predominantly expressed in the lateral preplacodal ectoderm (PPE), regulates the boundary between the NC and the placodes during the early steps of induction (Matsukawa et al., 2015). Specifically, Prdm12 inhibits the expression of *foxd3* and *sox8*, thereby restricting the NC domain during early neurula stages. Interestingly, in that study, Prdm12 did not have any inhibitory effect on Sox10 expression during the mid-neurula stages. Together with the results in mouse NC, this suggested that Prdm12 may not be solely antagonistic to NC development and/or may have stage-specific functions in the NC-GRN. Here, in *Xenopus laevis* embryos, we find that Prdm12 is expressed at low levels in the premigratory and early-migrating NC and is required for their EMT and migration, in a cell-autonomous manner. Our results further suggest that these regulations may not involve Prdm12’s histone methyl transferase activity. Downstream in Prdm12-dependent regulatory cascade, we uncover a regulatory connection between the Wnt signaling pathway and N-cadherin membrane localization in migratory NC cells. Together, our results depict a multifaceted role for Prdm12 during a key step of NC development and establishes a novel epistatic cascade linking signaling-mediated regulation and cellular migration.

## RESULTS

### Prdm12, a target of Pax3 regulates neural crest emigration in neurula stage embryos

In a previous transcriptomics study (Kotov et al., 2024) using a validated morpholino (MO) against Pax3 (Monsoro-Burq et al., 2005), we had identified Prdm12 as a potential target of Pax3 in the developing NC progenitors (Fig. S1A). We validated this observation in vivo by Pax3 knockdown and gain-of-function. At both early and late neurula stages, MO-mediated *pax3* depletion decreased *prdm12* expression, while *pax3* gain-of-function increased its expression both in the PPE and neural plate expression domains (Fig. S1B). As Pax3 is an essential regulator of the NC-GRN, we wondered whether Prdm12 played an overlooked role during NC development. Using two validated MOs (Thélie et al., 2015; Matsukawa et al., 2015), we find that indeed MO-mediated *prdm12* depletion decreases expression of the early pan-NC marker *snail2* during late-neurula stages (stage 17/18), with a stronger effect on the later pan-NC marker *sox10* (Figs. 1A-C, S2A-D). We validated this observation using the iNC assay, where co-injection of inducible forms of *pax3* and *zic1* into the early blastula ectoderm generates an enriched population of NC cells (Fig. 1D; Milet et al., 2013): At stage 18 equivalent, when NC EMT starts *in vivo*, *prdm12* depletion reduced expression of late-neurula-stage NC markers like *sox10*, although it did not significantly affect early-neurula-stage NC markers like *snail2* or *sox8*, uncovering a possibility that NC EMT, rather than induction, may be affected in the absence of *prdm12* (Fig. 1E). In the iNC assay, Prdm12 did not affect the expression levels of *twist1*, so we used *twist1* probe to assess NC migration *in vivo* by *in situ* hybridization (ISH). Prdm12 depletion leads to reduced NC migration streams in a MO concentration-dependent manner (Figs. 1F, G and S2E, F). Together, these observations introduced Prdm12 as an important regulator of NC development, especially during NC cells emigration.

**Fig. 1.**
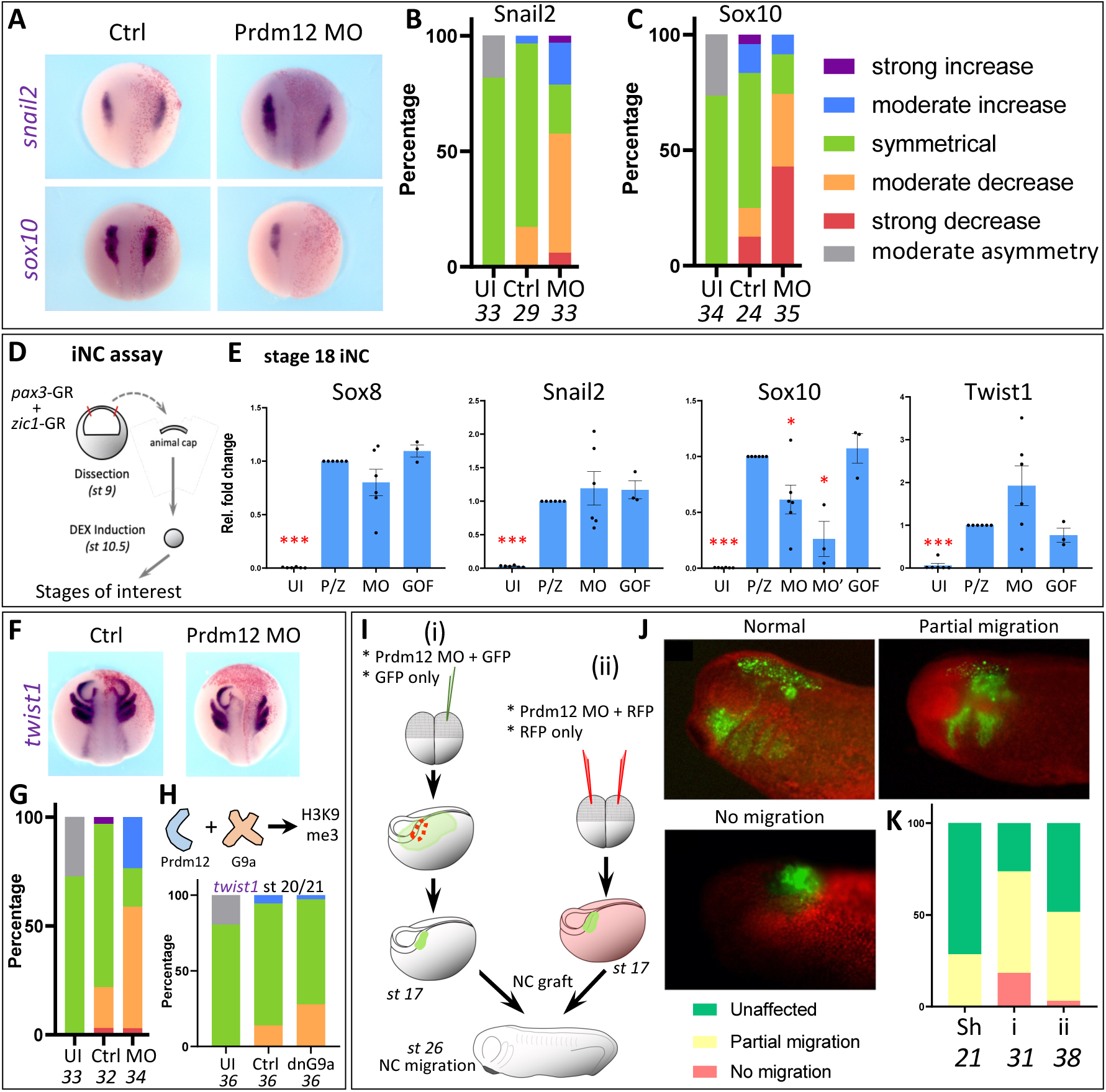
Prdm12 regulates the later stages of NC development. **(A)** ISH shows the effect of *prdm12* depletion (morpholino, MO) on *snail2* and *sox10* during late neurula (st 17/18) stage. **(B, C)** Quantification of phenotypes shown in (A) for *snail2* (B) and *sox10* (C). UI – uninjected, Ctrl – control, MO – *prdm12* knockdown. **(D)** Schematic representation of the different steps of the induced NC (iNC) assay. GR-constructs used here can be translocated to the nucleus by treatment with Dexamethasone. **(E)** RT-qPCR experiments depict the normalized relative expression of *sox8*, *snail2* and *sox10* in stage 18 iNC assay. UI – uninjected, P/Z – *pax3*-GR + *zic1*-GR, MO – P/Z + *prdm12* MO (20 ng), MO’ – P/Z + *prdm12* MO (40 ng), GOF – P/Z + *prdm12* mRNA. All conditions normalized and compared to P/Z. N ≥ 3. **(F)** ISH shows the effect of *prdm12* depletion on early NC migration (*twist1*, st 20/21). **(G)** Quantification of phenotypes shown in (E). UI – uninjected, Ctrl – control, MO – *prdm12* knockdown. Legend as in (B, C). **(H)** Prdm12 requires G9a to perform its transcriptional repressor function (Yang and Shinkai, 2013). ISH against *twist1* shows the effect of a dominant-negative version of G9a. Quantification of embryonic phenotypes. UI – uninjected, Ctrl – control, dnG9a – dominant-negative G9a. Legend as in (B, C). **(I)** Scheme of orthotopic NC graft: i – a *prdm12* depleted NC was grafted onto a control embryo, ii – a control NC was grafted onto a *prdm12* depleted embryo. **(J, K)** Depiction (J) and quantification (K) of phenotypes observed. Sh – control NC graft onto control embryo, i,ii – same as in I. In A, F, red staining depicts the injected side. Numbers under graphs depict number of embryos for each condition. Error bars depict standard error of mean. The significance of differences was calculated using Welch’s t-test (two-tailed); * depicts *p* < 0.01, *** depicts *p* < 0.0001.

From a biochemical perspective, Prdm12 is known to require its partner, G9a (or EHMT2), to induce methylation of histone proteins (Yang and Shinkai, 2013). When we introduced a dominant-negative G9a in the embryo, to abolish the epigenetic activity of Prdm12, we did not see a similar degree of NC migration defects (Fig. 1H). Therefore, the effect of Prdm12 on NC migration does not seem to exclusively depend upon its transcriptional repressor activity. This is in line with a recent study also showing non-enzymatic functions of Prdm12 during the development of the nociceptive sensory neurons (Tsimpos et al., 2024). Together, this strongly suggested that Prdm12 may display alternative functions in the NC-GRN.

Previous reports have shown that NC migration can be controlled cell non-autonomously by factors from the adjacent PPE, such as the matrix metalloproteinase MMP28 and the chemokine Sdf1, which are required for NC EMT and migration respectively (Escot et al., 2013; Gouignard et al., 2023; Theveneau et al., 2013). Since *Prdm12* depletion also affects PPE development (Fig. S2G, H; Matsukawa et al., 2015), which could in turn cause the defective NC migration, we assessed whether Prdm12 effect on NC migration was cell-autonomous or non-cell-autonomous by two types of orthotopic grafts. Firstly, to detect non-cell-autonomous effects, we grafted a lateral PPE explant onto a stage-matched host early neurula (stage 14) (Fig. S2I). At early tailbud stage (stage 26), we assessed NC migration through *twist1* expression. When a *prdm12* morphant PPE was grafted onto a wild type host, there were no significant effects on NC migration (Fig. S2J, K). Secondly, we checked cell-autonomous effects by grafting a NC explant onto a host neurula (Fig. 1I-K). A *prdm12* morphant NC grafted onto a wild-type host (Fig. 1Ii) showed strong defects compared to a wild-type NC grafted onto a *prdm12* morphant embryo (Fig. 1Iii). Collectively, these experiments suggested that Prdm12 mainly acts on NC migration cell-autonomously.

### Prdm12 is expressed at low levels in the neural crest

While our observations strongly suggested that Prdm12 regulates NC emigration cell-autonomously, no previous study had reported whether it is expressed in NC cells themselves. During neurula stages, *prdm12* is expressed in the lateral PPE and in two bilateral longitudinal stripes in the posterior neural plate (Matsukawa et al., 2015; Thélie et al., 2015). First, we confirmed the expression patterns of *prdm12* in *Xenopus laevis* embryos, using *in situ* hybridization (Fig. 2A). Similar to previous observations, at late gastrula stage 12.5/13, *prdm12* is expressed at the antero-lateral sides of the embryo. As neurulation proceeds, *prdm12* also starts to be expressed in two bilateral stripes in the neural plate. At tailbud stages 24/25, *prdm12* is expressed in the otic placodes, parts of the adenohypophyseal and trigeminal placodes, and in the neural tube. To compare the *prdm12* expression domain with the NC region, we examined the expression of *prdm12* with respect to *sox10*. At stage 17/18, although *prdm12* and *sox10* are not fully co-expressed in the same tissues, there is a small overlap between their expression domains (Fig. 2B, red arrowhead). Similarly, we observed a possible overlap at tailbud stages 24/25 (Fig. 2B). To validate these observations, we performed a set of microdissection experiments, followed by qRT-PCR to quantify *prdm12* gene expression. Firstly, we micro-dissected the anterolateral region of a late neurula stage embryo and divided it into 4 parts: the NC (1a), the ectoderm overlying the NC (1b), the lateral PPE (2) and the anterior PPE (3) (Fig. 1C). As expected, the lateral PPE exhibited the highest levels of *prdm12* expression, with slightly lower levels in the anterior PPE (Fig. 1D). The NC explant also expressed *prdm12*, albeit at much lower levels than placodal tissues, but significantly higher than in control tissue (naïve ectoderm). Concomitantly, explanted NC tissues maintained expression of *prdm12* up to a time equivalent to stage 22 (∼6h, grown at RT, 18°C), when the NC cells are migratory *in vivo* (Fig. S3A). We also tested whether *prdm12* is induced in the iNC assay, and observed detectable levels of *prdm12* in the induced NC caps as early as stage 14, with significantly higher levels at stage 18 (Fig. S3B). Finally, we performed a spatial transcriptomics FISH experiment (MERSCOPE, Vizgen) and observed *prdm12* expression in the *sox10+* NC cells at neurula and early tailbud stages (Fig. 2E). Sørenson-Dice scores depict a low level of coexpression of *prdm12* and *sox10* at stage 19 (post EMT, premigratory), with higher levels at stages 20 and 22 (migratory), comparable to *prdm12*/*six1* coexpression levels (Fig. 2F). The low score at stage 18 may be due to an incomplete overlap of the expression domains, which seems to gradually increase (Fig. 2E, clear arrowhead). Interestingly, as a control of FISH quality, *prdm12* also shows a low level of coexpression with *sox2* (neural plate) and *krt12.4* (epidermis), although it does not overlap with *sox17* (endoderm) (Fig. S3C-H), indicating that *prdm12* expression is restricted to the ectoderm. Together, these experiments showed that in addition to the lateral PPE, *prdm12* is also expressed endogenously in the NC cells, in particular in the cells positioned at the lateral edge of the NC domain, prone to emigration. Consequently, this suggested its potential involvement in NC cell EMT and migration initiation.

**Fig. 2.**
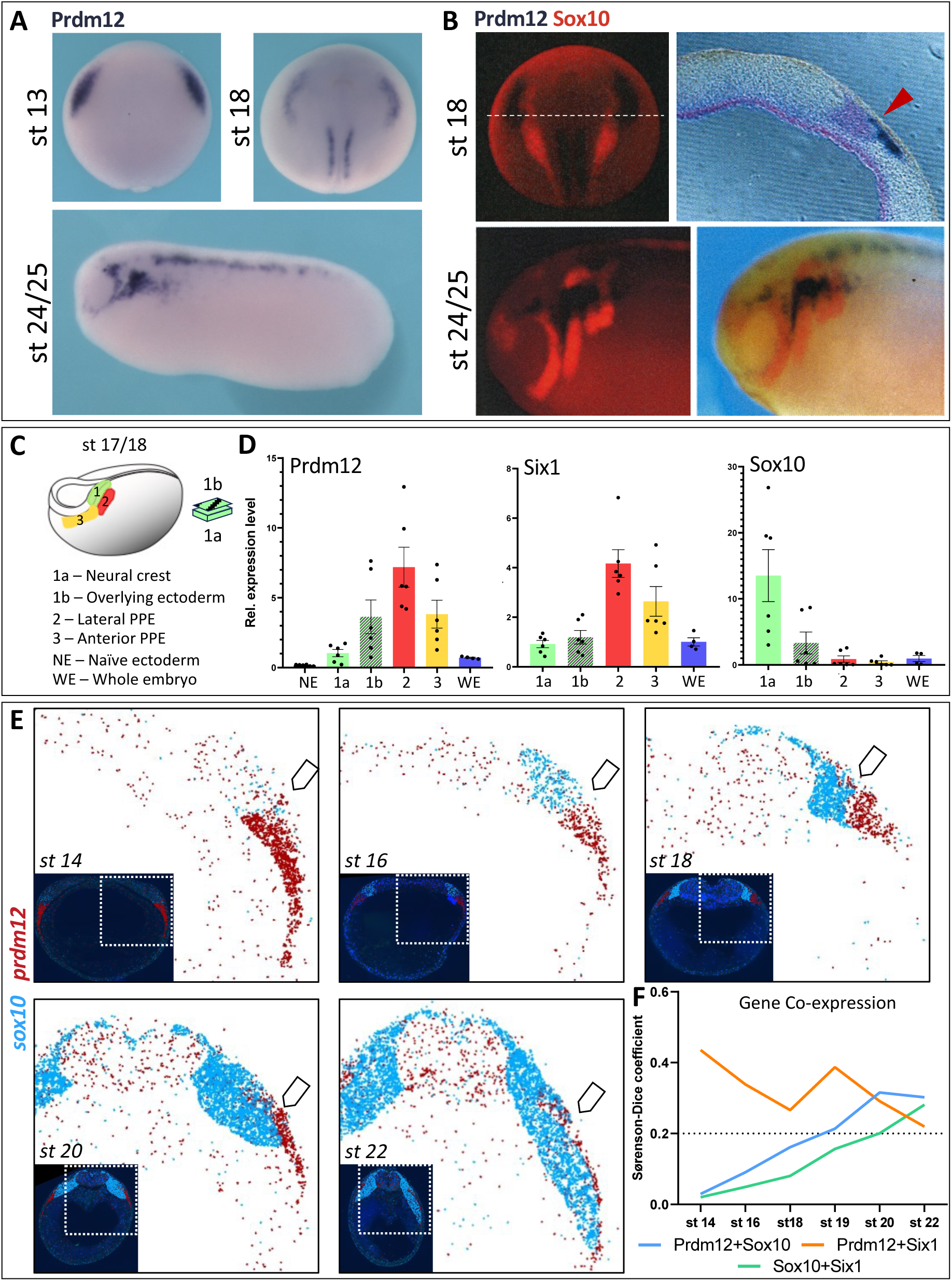
Prdm12 is expressed at low levels in the NC. **(A)** ISH shows the expression patterns of *prdm12* during early neurula (st 13, dorsal view), late neurula (st 18, dorsal view) and tailbud (st 24/25, lateral view) stages. *Prdm12* is expressed in the lateral preplacodal ectoderm (13 and 18), as lateral stripes in the posterior neural tube (18 and 24/25) and in the otic/adenohypophyseal placodes (24/25). **(B)** Double ISH shows the expression patterns of *prdm12* and *sox10* during late neurula (st 18) and tailbud (st 24/25) stages. White dotted line depicts plane of sectioning. Red arrowhead highlights the potential overlap. **(C)** Scheme of microdissection experiment at late neurula (st 17/18) stage. PPE – preplacodal ectoderm. **(D)** RT-qPCR experiments depict the normalized relative expression of *prdm12*, *six1* (PPE) and *sox10* (NC) in the tissues listed in (C). Naïve ectoderm is animal cap dissected at stage 9 and grown till stage 17/18. N = 6. **(E)** Spatial transcriptomics experiment depicts the expression patterns of *sox10* and *prdm12* in transverse sections of embryos at different stages. Main images depict the right dorsal side, inset images depict the whole section. Clear arrowhead depicts potential region of co-expression. **(F)** Sørenson-Dice coefficient depicts the degree of coexpression of different genes: x<0.2 – non-significant, 0.2<x<0.5 – moderate coexpression, 0.5<x – strong coexpression.

### Prdm12 regulates NC cell dispersion through N-cadherin function

Our experiments suggested that Prdm12 exerted cell-autonomous control over NC migration. To understand its molecular mechanism, we next looked at the cellular process of NC migration *in vitro* (Fig. 3A). In *Xenopus* embryos, the antagonistic mechanisms of contact inhibition of locomotion (CIL) and chemotactic co-attraction (CoA) ensure that the NC cells migrate as a tight group of cells (Carmona-Fontaine et al., 2008; Carmona-Fontaine et al., 2011). However, under *in vitro* conditions over a fibronectin substrate, NCCs tends to migrate as unattached single cells, which leads to dispersion of NC explants (Scarpa et al., 2015). In this experimental setup (Fig. 3A), depletion of *prdm12* significantly reduced the ability of the NC cells to disperse *in vitro* (Fig. 3B, C; Movie 1A, B). This dispersion defect could be due to two reasons: either the NC cells possessed a defective migration machinery (i.e., the cytoskeleton), or the NC cells were unable to detach from the cluster (i.e., cell adhesion defect). To explore these hypotheses, we injected embryos with minimal amounts of fluorescently tagged markers: SF9-mNeonGreen to detect myosin II (Arnold et al., 2019), LifeAct-mCherry to detect actin, and CAAX-miRFP670 to detect the cell membrane. Similar to control NC cells, *prdm12* morphant NC cells successfully attached to the substrate. However later on, when the control cells had lost their intercellular connections and become single cells, *prdm12* morphant cells remained clustered (Fig. 3D, E; Movie 2A, B). Importantly, morphant cells formed exploratory lamellipodial protrusions (Fig. 3D, yellow arrowheads) but did not detach to form single cells. We also observed sporadic reductions in myosin and actin levels. However, since this could be attributed to microinjection variations, we did not investigate this further.

**Fig. 3.**
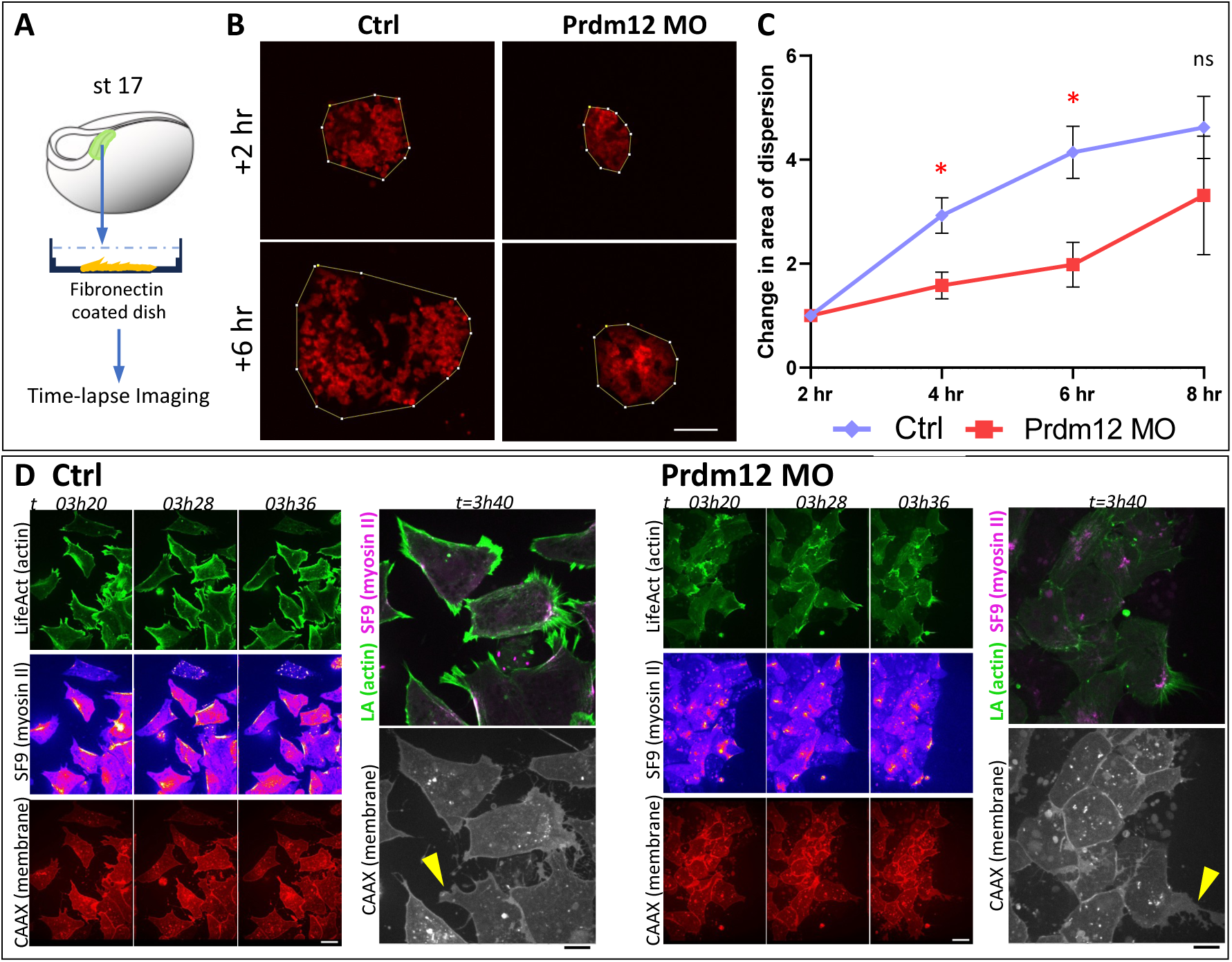
Prdm12 depletion perturbs neural crest dispersion during its migration. **(A)** Scheme of experiment to assess NC migration *in vitro*. NC explants were micro-dissected prior to EMT at stage 17 and placed on a fibronectin-coated surface. **(B)** GFP-labeled NC cells were allowed to migrate on fibronectin for 8 hours (t = 0 at time of dissection) at 18°C. Yellow line shows the total area covered by the explant. Scale bar, 100 μm. **(C)** Quantification of relative change in area over time for assay in B. Area at t = 2h set as 1. For control explants, cells migrated out of frame between 6h and 8h, leading to inaccurate measurements. N = 6 (ctrl) or 3 (MO). **(D)** Time-lapse images showing the migration of NC explants in control or *prdm12* knockdown conditions. The cytoskeleton in visualized by the fluorescence of live biomarkers – Lifeact for actin, Sf9 for myosin II and CAAX for membrane. White arrowheads depict myosin II localization, yellow arrowhead depicts lamellipodial formation in *prdm12* morphant NC cells. Scale bar, 10 μm. Error bars depict standard error of mean. The significance of differences was calculated using Welch’s t-test (two-tailed); * depicts *p* < 0.01, ns depicts *p* > 0.05.

Two previous reports have shown a similar dispersion defect phenotype. The first involves cadherin function. Contact-inhibition-of-locomotion (CIL) in the NC is dependent on the switch of expression from E-cadherin (*cdh1*) to N-cadherin (*cdh2*) (Scarpa et al., 2015). Consequently, a loss of N-cadherin leads to adhesion defects and the loss of CIL behaviour, and NC cells are unable to disperse. Interestingly, a high proportion of *prdm12* expressing *sox10*+ NC cells also expressed *cdh2* (Fig. 4A). Consequently, since *prdm12* depletion seemed to phenocopy the loss of N-cadherin, we investigated whether *prdm12* regulates the expression and/or localization of N-cadherin. In the iNC assay at stage 18, *prdm12* depletion led to significantly reduced levels of N-cadherin in a morpholino dose-dependent manner (Fig. 4B). To check whether this change in the transcript level also affects the N-cadherin protein and its localization, we performed immunofluorescence (IF) staining of *in vitro* migratory NC explants at t=6h30 after dissection (roughly equivalent to stage 22 *in vivo*, at 18°C). Since N-cadherin localizes to adherens junctions formed between two cells, we quantified N-cadherin localization in clustered NC cells (Fig. 4C, D). In control NC explants, N-cadherin localized to the intercellular junctions, with no significant differences between junctions formed between cells at the edge of the explant and junctions formed between interior cells. However, in *prdm12* morphant NC cells, the amount of N-cadherin localized to the intercellular junctions (both interior and edge) was significantly reduced. In certain cases, N-cadherin was almost completely absent near cell boundaries, which could also suggest defective intercellular adherens junctions. This reduction in N-cadherin upon *prdm12* depletion suggested a possible mechanism how Prdm12 may regulate NC migration. However, the introduction of chicken N-cadherin (Scarpa et al., 2015) was not sufficient to rescue the NC migration defects upon loss of *prdm12* (Fig. 4E, F), which suggested the presence of additional regulatory mechanisms. Since the expression of N-cadherin involves a switch from E-cadherin, we also looked at whether the levels and/or localization of E-cadherin is affected upon *prdm12* depletion. Contrary to one previous study (Scarpa et al., 2015) while supporting another (Huang et al., 2016), we observed E-cadherin expression in the migratory NC, at t=6h30 (∼stage 22, at 18°C) post dissection (Fig. 4G). However, Prdm12 did not seem to regulate the levels or localization of E-cadherin in the NC (Fig. 4H).

**Fig. 4.**
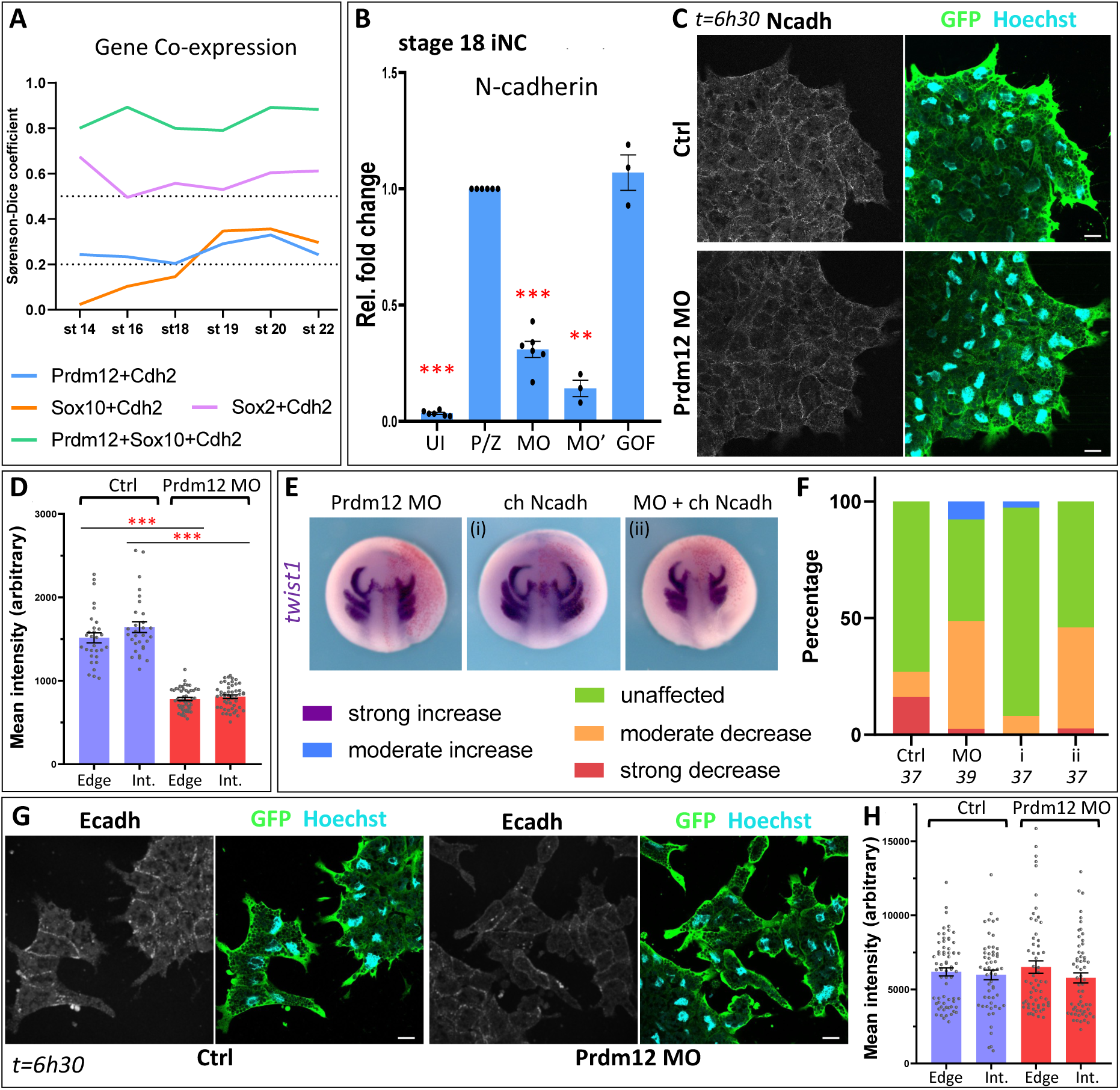
Prdm12 depletion reduces the level of N-cadherin at intercellular junctions. **(A)** Sørenson-Dice coefficients calculated from the spatial transcriptomics experiment: x<0.2 – non-significant, 0.2<x<0.5 – moderate coexpression, 0.5<x – strong coexpression. N-cadherin shows low levels of co-expression with either *prdm12* or *sox10* around migration stages. However, a significant proportion of *prdm12*/*sox10* double-positive cells show expression of N-cadherin. As per previous reports, N-cadherin consistently shows a high overlap with *sox2* expression. **(B)** RT-qPCR experiments show the effects of *prdm12* depletion on the expression of N-cadherin (*cdh2*) in stage 18 iNC assay. UI – uninjected, P/Z – *pax3*-GR + *zic1*-GR, MO – P/Z + *prdm12* MO (20 ng), MO’ – P/Z + *prdm12* MO (40 ng), GOF – P/Z + *prdm12* mRNA. All conditions normalized and compared to P/Z. N ≥ 3. **(C)** IF images showing N-cadherin expression in NC explants on fibronectin, 6h30 (at 18°C) after dissection in control or *prdm12* knockdown conditions. GFP labels injected NC cells. Scale bar, 15 μm. **(D)** Quantification of mean intensity of N-cadherin at intercellular junctions between cells at the edge or between interior cells. n = 10 cells, N = 3 (Ctrl) or 5 (MO) explants. **(E)** ISH analysis depicts the effect of introducing chicken N-cadherin in *prdm12* morphant NC. Red staining depicts the injected side. **(F)** Quantification of phenotypes shown in (E). Numbers under graphs depict number of embryos for each condition. Ctrl – control, MO – *prdm12* knockdown, i – chicken N-cadherin, ii – *prdm12* MO + chicken N-cadherin. **(G)** IF images showing E-cadherin (*cdh1*) expression in NC explants on fibronectin, 6h30 (at 18°C) after dissection in control or *prdm12* knockdown conditions. GFP labels injected NC cells. Scale bar, 15 μm. **(H)** Quantification of mean intensity of N-cadherin at intercellular junctions between cells at the edge or between interior cells. n = 10 cells, N = 5 explants. Error bars depict standard error of mean (SEM). The significance of differences was calculated using Welch’s t-test (two-tailed); ** depicts *p* < 0.001, *** depicts *p* < 0.0001.

### Prdm12 controls NC migration through regulating the non-canonical (PCP) Wnt pathway

Wnt signaling plays an important role not only during NC induction, but also during EMT and the initiation of NC emigration. Interestingly, the regulation of both canonical and non-canonical (planar cell polarity, PCP) Wnt signaling in *Xenopus* is not binary and precise levels are required for successful migration (De Calisto et al., 2005; Maj et al., 2016). Interestingly, a dysregulation of canonical Wnt signaling output leaves NC cells unable to detach, a phenotype similar to that of *prdm12* depletion. This prompted us to investigate whether Prdm12 affects either branch of the Wnt signaling pathway. First, in the iNC assay at stages 16 and 18, possibly when the mRNA of genes required for EMT and migration are transcribed, we observed that *prdm12* regulates the expression of multiple Wnt components. *Prdm12* depletion led to significantly decreased expression of *axin2* (canonical Wnt target gene) and *ror2* (non-canonical Wnt co-receptor), while it increased the expression of *gsk3a* (canonical Wnt antagonist), *dvl2* and *fzd7* (genes involved in both pathways) (Fig. 5A). A decrease in *axin2* and an increase in *gsk3a* suggested a reduced level of canonical Wnt signaling, while a decrease in *ror2* levels also suggested a concomitant reduction in non-canonical Wnt signaling. Next, we wanted to check whether the changes in gene expression also had a concomitant effect on the signaling output. We used two luciferase reporter assays: a Tcf3-based TOPFLASH assay for canonical Wnt (Promega) and an ATF2-based assay for non-canonical Wnt (Ohkawara and Niehrs, 2011). We introduced a small amount of Wnt activators in the naïve ectoderm (animal cap) to raise the endogenous activity levels and subsequently checked the effect of *prdm12* depletion to depict whether Prdm12 had a direct effect on the respective pathways. We observed that *prdm12* depletion led to reduced signaling outputs of both the canonical and the non-canonical Wnt pathways (Figs. 5B and S4A). Interestingly, similar to the effect on NC migration, it seemed this perturbation was only weakly dependent upon the transcriptional repressor activity of Prdm12: Introduction of the dominant-negative G9a construct in this assay did not lead to a similar significant decrease, except in the case of Dvl2 ΔDIX (canonical Wnt activity). Nonetheless, this confirmed that Prdm12 regulated both the arms of Wnt signaling, albeit through its non-enzymatic activity.

**Fig. 5.**
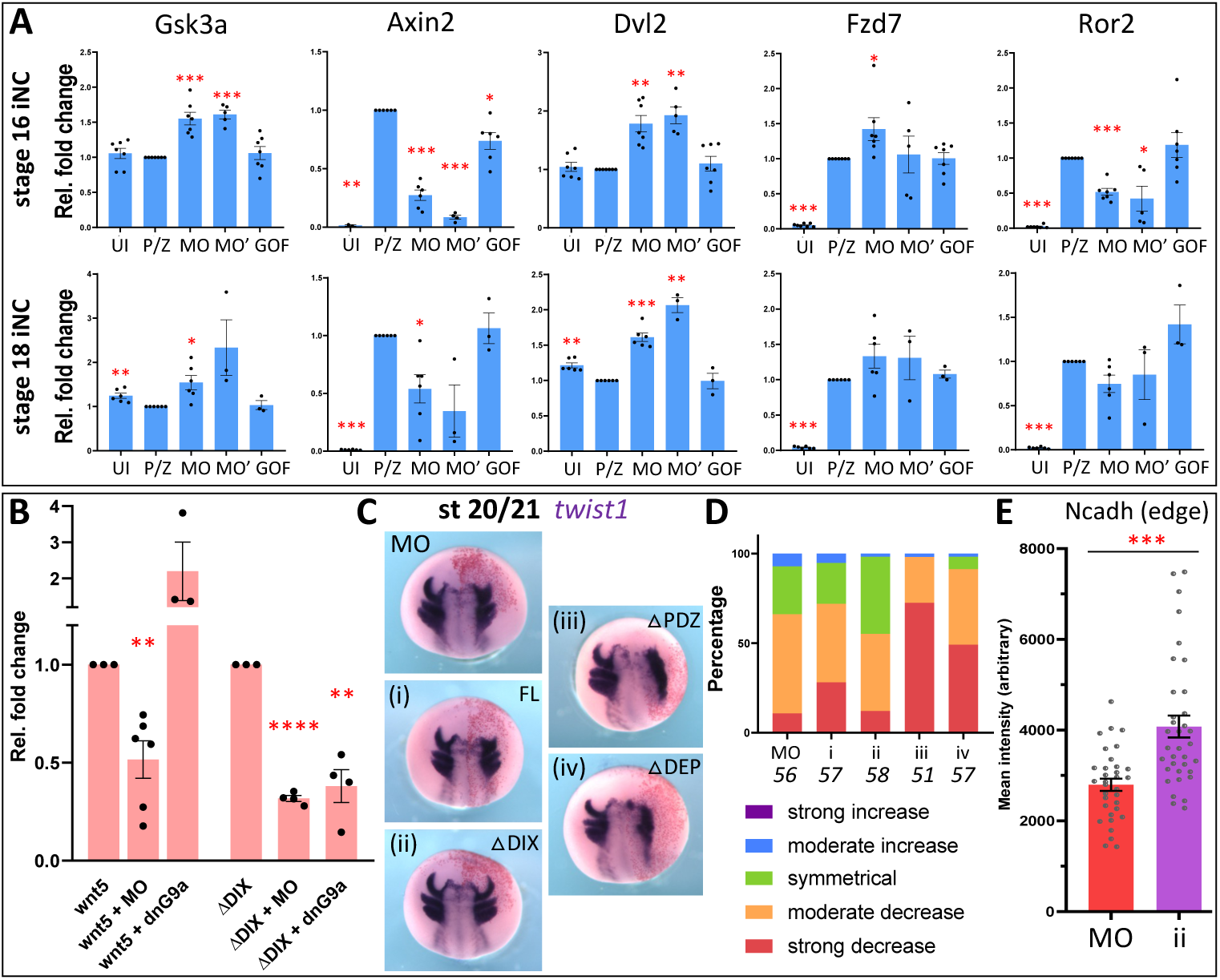
Prdm12 regulates N-cadherin through ncWnt signaling. **(A)** RT-qPCR experiments show the effects of *prdm12* depletion on the expression of Wnt components in stage 16 or 18 iNC assay. MO – P/Z + *prdm12* MO (20 ng), MO’ – P/Z + *prdm12* MO (40 ng), GOF – P/Z + *prdm12* mRNA. All conditions normalized and compared to stage-specific P/Z. N ≥ 3. For *axin2* stage 16, N = 2 for UI. **(B)** ATF2-Luciferase assay shows the effect of *prdm12* depletion on non-canonical Wnt signaling readouts in animal cap explants microdissected at stage 9 and grown till stages 17/18. MO – *prdm12* knockdown, ΔDIX – Dvl2 without the DIX domain, dnG9a – dominant-negative G9a. N ≥ 3. **(C)** ISH shows the effect of Dvl2 in *prdm12* depleted NC (*twist1*, st 20/21). MO – *prdm12* knockdown, i – MO + *dvl2*, ii – MO + *dvl2* ΔDIX, iii – MO + *dvl2* ΔPDZ, iv – MO + *dvl2* ΔDEP. Δ depicts domain deletion. Red staining depicts the injected side. **(D)** Quantification for phenotypes shown in (C). Numbers under graphs depict number of embryos for each condition. **(E)** Quantification of mean intensity of N-cadherin at intercellular junctions between cells at the edge in morphant (MO) or morphant + *dvl2* ΔDIX (ii) NC explants. n = 10 cells, N = 5 explants. Error bars depict standard error of mean. The significance of differences was calculated using Welch’s t-test (two-tailed); *** depicts *p* < 0.0001.

From the combination of genes affected, it was difficult to predict which arm of Wnt signaling was more important in the context of Prdm12 and NC migration. To differentiate between the effects of the two arms, we attempted to rescue the NC migration defects upon *prdm12* depletion with different constructs of *dvl2*. Dvl2 plays important roles in both the canonical and non-canonical Wnt pathways, and interacts with different components in a domain-dependent manner (Kratzer et al., 2020). For its functions in the canonical Wnt pathway, Dvl2 requires its N-terminal DIX domain, while the DEP domain is important for its role in the non-canonical Wnt pathway (Sharma et al., 2018; Shi, 2020). The PDZ domain is required for both arms, and a few studies have shown that the DEP domain is also important for canonical signaling in *in vitro* conditions (Gammons et al., 2016; Paclíková et al., 2017). We co-injected the Prdm12 morpholino with low amounts of *dvl2* FL (full length), *dvl2* ΔDIX (*dvl2* lacking the DIX domain), *dvl2* ΔPDZ (*dvl2* lacking the PDZ domain) or *dvl2* ΔDEP (*dvl2* lacking the DEP domain). Co-injection with *dvl2* FL was not able to rescue the migration defects, while we observed a partial rescue of the migration defects in case of *dvl2* ΔDIX (Fig. 5C, D). On the other hand, the migration defects worsened when Prdm12 MO was co-injected with either *dvl2* ΔPDZ or *dvl2* ΔDEP (Fig. 5Ciii, Civ). Further, when we attempted to rescue migration defects through modulating the canonical Wnt pathway directly (Maj et al., 2016), we did not observe any significant rescue, but rather a worsening of the defect phenotype (Fig. S4B). Thus, the regulation of the non-canonical Wnt (PCP) pathway by Prdm12 seemed more crucial during NC migration.

The non-canonical Wnt pathway has been shown to regulate structural components of the cytoskeleton directly. Further, Dvl2 directly regulates the dynamics of Cdh3 (Huebner and Wallingford, 2022). In *Xenopus* NC, non-canonical (PCP) Wnt is also required for contact-inhibition-of-locomotion, which is mediated by N-cadherin (Carmona-Fontaine et al., 2008). Thus, we checked whether *dvl2* ΔDIX could rescue the localization of N-cadherin. Indeed, introduction of *dvl2* ΔDIX in a *prdm12* morphant background significantly rescued the membrane localization of N-cadherin (Fig. 5E). Thus, this uncovers an epistatic relationship where Prdm12 regulates the non-canonical Wnt (PCP) pathway, which then controls the localization of N-cadherin. Together, this suggests a novel mechanism by which Prdm12 regulates NC migration.

## Discussion

The NC-GRN is one of the most studied in vertebrate systems, with studies focusing on all the steps of its early development. Over the last few decades, the GRN has grown considerably, with added genes including transcription factors, signaling components and epigenetic modifiers. However, a broader connection between different aspects of regulation, for example between signaling and cellular mechanisms, still remains missing. In such a case, it becomes imperative to revisit some of these factors, with the motive of understanding their molecular mechanisms and the possible cooperation between them. In this study, we adopted a more candidate-based genetics approach: we focused on a particular gene and elucidated its molecular function. While doing so, we uncovered a novel axis of regulation, which links signaling and cellular migration.

### Prdm12 is required to prime the Xenopus NC for migration

The PRDM family of proteins are PR (positive-regulatory) domain-containing proteins that display histone methyl transferase activity (Casamassimi et al., 2020). Most of these genes are implicated in cancer, while a few (Prdm1, Prdm6) are also known to be important for NC development (Hong et al., 2022; Powell et al., 2013; Prajapati et al., 2019). In *Xenopus laevis*, Prdm12 is expressed in the lateral PPE and the p1 progenitors of the neural tube (Matsukawa et al., 2015; Thélie et al., 2015), and is important for neurogenesis of nociceptors. Here, we demonstrate that Prdm12 is an important regulator of NC EMT and migration in *Xenopus laevis*. A previous report in the same system had shown that placodal expression of Prdm12 restricts the NC boundary during early neurula stages (Matsukawa et al., 2015). This occurs through the transcriptional repression of the early NC markers like *foxd3* and *sox8* (Fig. 6, Induction stage). Here, we show that Prdm12 is co-expressed with NC markers, with increased expression levels as neurulation proceeds, with higher levels in the NC during EMT and later stages and that Prdm12 exerts cell-autonomous control over NC EMT and migration (Fig. 6, EMT – zone iii). Depletion of *prdm12* leads to a significant decrease only in the NC late gene *sox10*, but not other EMT markers. *Prdm12*-deficient NC cells do exhibit partial migration, which could suggest that EMT, although perturbed is not completely abrogated. Sox10, a transcription factor, is not only required for NC-EMT and migration (Honoré et al., 2003; McKeown et al., 2005; Schock and LaBonne, 2020), but also regulates the specification and development of NC derivatives such as melanocytes, glia, neurons and several others (Harris et al., 2013; Kellerer et al., 2006). Interestingly, loss of PRDM12 causes a reduction in the number of SOX10+ NC cells in mouse embryos at stages E11.5 and E12.5, hinting at a possible conserved regulatory mechanism (Bartesaghi et al., 2019). However, while extensive transcriptomic studies have revealed genes downstream of *sox10* in NC derivatives (Fufa et al., 2015; Girard and Goossens, 2006), we still lack a comprehensive list of *sox10* targets during neurula stages, which forms an interesting arena for further research. Potentially, this could uncover yet another axis through which Prdm12 may regulate NC-EMT.

**Fig. 6.**
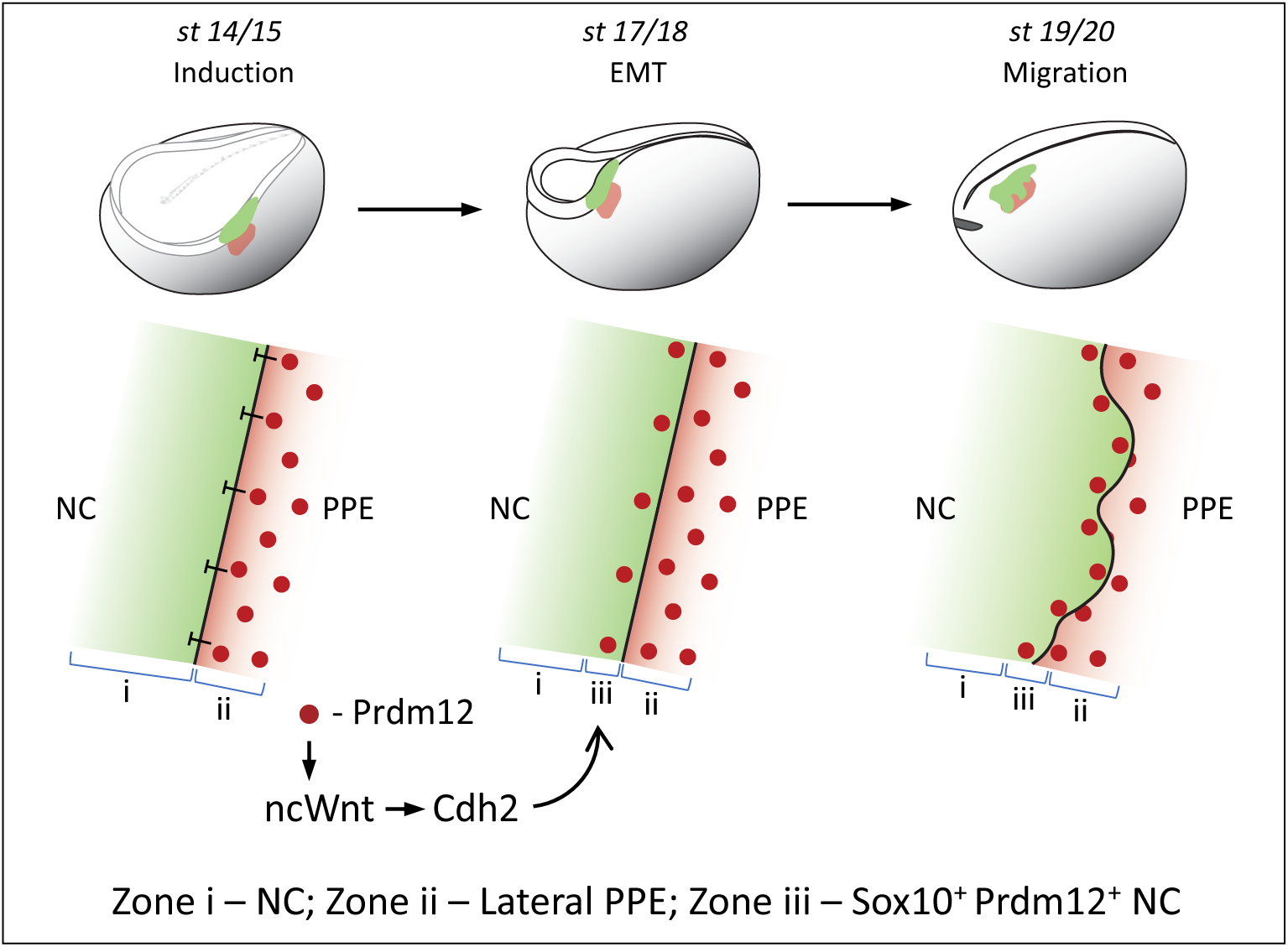
A novel function for Prdm12 during NC migration in *Xenopus laevis*. At early-mid neurula stages, Prdm12 inhibits the expression of early NC genes *foxd3* and *sox8*, thereby establishing the boundary between the NC and the lateral preplacodal Ectoderm (PPE). At late neurula stages, the lateral-most NC cells express low levels of *prdm12*. These *sox10*/*prdm12* double-positive cells express N-cadherin (*cdh2*), induced by Prdm12 through the non-canonical Wnt (ncWnt) signaling pathway. Subsequently, these NC cells might be the first ones to start migration.

There is still a debate whether the lateral-most NC cells in *Xenopus* migrate first, as in zebrafish and chick trunk (Richardson et al., 2016; Theveneau and Mayor, 2012a; Theveneau and Mayor, 2012b). All the NC cells exhibit similar migratory capacities, although due to space constraints and contact-inhibition-of-locomotion, the lateral-most NC cells migrate first (Alfandari et al., 2010; Barriga and Theveneau, 2020; Carmona-Fontaine et al., 2008; Marchant et al., 2022). However, in a previous single-cell transcriptome study, we have shown that the premigratory NC cells display extensive transcriptomic heterogeneity (Kotov et al., 2024). In this study, *prdm12* seems restricted to the lateral edge of the NC tissue, which we assume suggests different characteristics of the NC cells depending upon position.

Concomitantly, from our spatial transcriptomics data, we did observe a significant enrichment of N-cadherin in *sox10*-positive cells also expressing *prdm12*. N-cadherin is a necessary marker of the mesenchymal nature of post-EMT NC cells, contributing both to their delamination and migratory capabilities (Bahm et al., 2017; Scarpa et al., 2015). That we observe Prdm12 is required for N-cadherin expression, further strengthens our assumption. We hypothesize that *prdm12* is co-expressed in a small cohort of *sox10*+ NC cells at the lateral edge, which then express N-cadherin and undergo delamination and migration (Fig. 6, Migration – zone iii). It would be interesting to see if *sox10* also regulates the expression of N-cadherin. This could either be direct or indirect; Sox10 is known to interact with Twist1 (in the cardiac NC), a gene which can directly upregulate N-cadherin expression (Alexander et al., 2006; Vincentz et al., 2013; Yang et al., 2004). Thus, Sox10 and Prdm12 together may impart special characteristics to a small cohort of NC cells, which may then delaminate and/or migrate in a temporally-controlled manner. Almost all previous studies have used a smaller piece of the NC for *in vitro* analysis, thereby losing the spatial variation. In the future, for validation, it would be interesting to see if cells from an entire NC explant start to migrate dependent upon their position in the embryo.

### Prdm12 functions through the non-canonical (PCP) Wnt pathway

Signaling through the non-canonical (planar cell polarity/PCP) Wnt pathway, mediated by Wnt11, Wnt11r, Fzd7, Ror2 and Par3, is required for NC migration (De Calisto et al., 2005; Matthews et al., 2008; Moore et al., 2013; Podleschny et al., 2015). Prdm12 regulates the signaling output of this pathway, probably through its regulation of factors like Fzd7 and Ror2. Interestingly, previous transcriptomic studies have shown that Prdm12 is required for the expression of Wnt11 (non-canonical Wnt ligand), while it represses genes like integrins (functions in cell-ECM contact) and Wnt8a (canonical Wnt ligand) (Landy et al., 2021; Thélie et al., 2015). In the NC, the PCP pathway plays an important role in mediating contact-inhibition-of-locomotion, which also requires N-cadherin. The PCP pathway is known to interact with cadherins: both E and N-cadherin interact with small GTPases, which then control cell polarity and transcription/cell cycle (Charrasse et al., 2002; Du et al., 2014; Ouyang et al., 2013). Further, while Vangl2, a core PCP protein, regulates E-cadherin and cooperates with N-cadherin for cellular polarization (Nagaoka et al., 2014; Nagaoka et al., 2023), JNK activity, which is activated by PCP signaling, regulates the expression of both E- and N-cadherin (Niu et al., 2016; You et al., 2013). We show here that the non-canonical activity of Dvl2 rescues the localization defects of N-cadherin. However, in the context of the NC, it remains to be seen which component of the PCP signaling is responsible for this process.

### Non-epigenetic functions of Prdm12 – possible protein-protein interactions

Like the other members of its family, Prdm12 contains a PR domain and 3 zinc finger motifs, and acts a histone methyl transferase. However, Prdm12 does not possess an intrinsic enzymatic activity, but rather functions by recruiting its partner EHMT2/G9a through its zinc-finger domains (Yang and Shinkai, 2013). Prdm12 also interacts with Bhlhe22, which imparts it both activating and inhibiting functions (Yildiz et al., 2019). However, the epigenetic activity of Prdm12 does not account for all of its functions. A recent report showed that Prdm12’s interaction with G9a is dispensable for its functions in neurogenesis, indicating possible non-enzymatic roles of Prdm12 (Tsimpos et al., 2024). Similarly, here we see that the effect of Prdm12 on the non-canonical Wnt pathway does not require its enzymatic function. Based on the presence of zinc-finger domains, we can hypothesize that Prdm12 may also possess protein-binding functions, which influences the function of other factors. In the future, a dedicated search for such partners may uncover further novel roles of this molecule.

Altogether, our work uncovers the possible mechanisms by which Prdm12 regulates NC EMT and migration. This establishes a novel epistasis relationship in the context of the *Xenopus* NC, strengthening the NC-GRN and generating a link between signaling and migration. Our findings also display Prdm12’s multifaceted roles, potentially via protein-protein interactions, opening avenues for exploring its non-epigenetic functions. Concomitantly, it remains to be seen whether these modes of regulation are independent of each other, or if they cooperate.

## Materials and methods

### Animal handling and developmental staging

Embryos were obtained from *Xenopus laevis* by *in vitro* fertilization using standard methods described in (Sive et al., 2000). Embryos were staged according to (Nieuwkoop and Faber, 1994). Animal use followed recommendations of the European Community (2010/63/UE) and international guidelines (Authorization APAFiS#36928-2022042212033387-v1).

### Microinjections, constructs and reagents

Embryos were injected at different blastula stages for different experiments: at 2-cell stage (for all iNC assays), at 4-cell stage (for all Prdm12 genetic manipulations) and at 8-cell stage (for all rescue experiments). For Prdm12 depletion, we used two previously validated morpholinos (GeneTools): MO1 – 5’-GCAGCACCGAGCCCATCATTAATTC-3’ (Matsukawa et al., 2015) and MO2 – 5ʹ-CATTAATTCTGCCTGCGAGTCTGAC-3ʹ (Thélie et al., 2015). After comparing MO1 and MO2, we observed similar effects with respect to NC induction and migration (Fig. S2A-F). Thus, we used MO2 for all subsequent experiments. Control (Ctrl) depicts the use of either the standard control MO 5’-CCTCTTACCTCAGTTACAATTTATA-3’ (Gene Tools) or just ß-gal. For labelling injected sides, a co-injection of either ß-gal or GFP was used. The doses and the type of microinjection used for each assay is indicated in Table S1.

### Whole embryo in situ hybridization

Whole-mount embryo *in situ* hybridization was done as in (Monsoro-Burq, 2007). DIG-labeled or Fluorescein-labeled antisense probes were synthesized from plasmid templates. After staining, whole embryos were imaged using Zeiss Lumar equipped with an Icc colour camera.

### Tissue explants, iNC (induced neural crest) assay and grafting

The NC (or its surrounding tissues) was microdissected from stage 14, stage 17/18 or stage 20 embryos (Plouhinec et al., 2014; Kotov et al., 2024). For RT-qPCR experiments (Fig. 2C, D), 6 biological replicates were done, with each sample gathering 3 identical tissue explants from 3 sibling embryos. For imaging NC *in vitro* (Figs. 3, 4; Milet and Monsoro-Burq, 2014), each explant was analyzed individually. Post dissection, the NC explant was placed on a fibronectin-coated dish. A small piece of coverslip with edges dipped in non-reactive grease was placed for 15 minutes to facilitate attachment. The co-injection of the NB transcription factors Pax3 and Zic1 into the animal cap (naïve ectoderm) recapitulates early NC development, a minimal system to study NC called “induced neural crest” assay (iNC; Milet et al., 2013). To produce iNCs, both cells of 2-cell stage embryos were co-injected with hormone-inducible Pax3 and Zic1, along with other reagents as needed for each experiment (Table S1). Grafting of NC was carried out as in (Milet and Monsoro-Burq, 2014). A similar technique was used to perform grafting of the lateral preplacodal ectoderm (Figs. 1 and S2).

### Neural crest dispersion assay and live imaging

The microdissected NC (either whole or a ∼1/6th fragment) was placed on a fibronectin-coated Fluorodish (40 µg/ml in ¾ NAM with gentamycin). A piece of coverslip was gently placed on top of the explant for the first 30-45 minutes, to facilitate attachment prior to start of live imaging. For the NC dispersion assay (Fig. 3A-C), one image was collected every 30 minutes, for 10 hours. Cytoskeleton dynamics (Fig. 3D) was assessed using tagged Lifeact (for actin) and sf9 (for myosin II), imaged every 15 seconds for 20 minutes. All explants were imaged using a Nikon Eclipse Ti microscope equipped with a Yokogawa spinning disk and Photometrics sCMOS camera. All steps were conducted at RT (18°C).

### RT-qPCR

Samples collected in 50 µl of medium (1/3 MMR or 3/4 NAM) were lysed in 400 µl of Trizol (Invitrogen). Total RNA was extracted according to the manufacturer’s instructions, and 250-500 ng was used to prepare cDNA using random hexamers and MMLV reverse transcriptase (Promega). RT-qPCR was done with 5X SYBR mastermix (Biorad). All gene expressions were normalized against *odc* housekeeping gene. For each experimental sample in iNC assay, normalized gene expressions were compared to those in control iNCs (pax3-GR and zic1-GR injected). For other experiments, expressions were compared to a standard loading control (an equal proportion cDNA mix from whole embryos ranging from stage 9 to stage 22). Primers are shown in Table S2.

### Immunofluorescence, imaging, and quantification of cadherins

Explants were fixed (3.7% formaldehyde in ¾ NAM for 30 minutes), washed in PBS, permeabilized with 0.1% Triton-X in PBS, blocked in 5% BSA-3% FBS in PBS, and incubated with primary antibodies overnight at 4°C and secondary antibodies for 2h at RT. Primary antibodies include anti-N-cadherin (MNCD-2, DSHB, 1:50), anti-E-cadherin (BD610181, BD Biosciences, 1:500), anti-GFP (GFP-1020, aveslabs, 1:1000), and secondary antibodies included Goat anti-rat IgG AlexaFluor 647, Goat anti-mouse IgG2a AlexaFluor 555, Goat anti-chicken IgY AlexaFluor 488 (Invitrogen, all at 1:1000). Nuclei were stained by Hoechst 33342 (0.2 µg/ml, Thermo Fisher). Explants were imaged on a Leica SP8 microscope. Image analysis was done using Fiji. For quantifying N-cadherin and E-cadherin expression, a spline-fit multi-point line of thickness 10 was drawn across the junction and the mean intensity was plotted.

### Luciferase assay

The level of Wnt signaling was assessed through the TOPFLASH construct (for canonical Wnt, Promega) or ATF2-luciferase (for non-canonical Wnt; (Ohkawara and Niehrs, 2011). TOPFLASH or ATF2-luciferase plasmid (25 pg) was co-injected with Renilla luciferase plasmid (5 pg, internal control), together with other reagents (Table S1). Animal caps were dissected at stage 9, grown until the stage of interest. For each condition, 3 animal caps were pooled into 30 µl 1X Passive lysis buffer. Firefly luciferase activity was measured using a Dual-Luciferase assay system (Promega) in a Mithras LB 940 plate reader (Berthold Technologies), and normalized to the level of Renilla luciferase.

### Statistical analysis and data representation

For comparison, we used Student’s t test with Welch’s correction to compare datasets. The statistical tests and all graphs were realized using GraphPad Prism 8.

### Spatial transcriptomics (multiplex-FISH by MERSCOPE)

*Xenopus.laevis* embryos were fixed, dehydrated and embedded in paraffin and cut as 7 μm sections. The sections were mounted on Vizgen MERSCOPE slides and dried overnight at 42 ℃. Subsequently, tissue clearing and multiplex FISH imaging steps were performed following the Vizgen instruction manual, with small adaptations. In short, a cell boundary staining was applied, followed by hybridization with the custom-made gene panel probes. Slides were imaged with MERSCOPE equipment, and processed with the software version 233. At least 3 sections from 3 different embryos were used for each stage. The cells were resegmented using Vizgen post-processing tools (VPT version 3.1) with an in-house trained Cellpose2 model. Transcripts were then assigned to the new cell segmentation. Transcript ≤ 1 count in a cell was regarded as background, cells with a total count < 6, a volume > 20000 μm^3^, and number of genes detected < 15 were removed. For co-expression analysis, we regarded positive cells as counts ≥ 3 detected for each gene. Sørensen-Dice coefficient was counted using the total number of cells per stage expressing each gene as follows: 2(A ∩ B) / (A + B), where A and B depict genes A and B. For *prdm12*+*sox10*+*cdh2* (Fig. 4A), *prdm12*+*sox10* was considered as set A and *cdh2* was considered as set B.

---

**Fig. S1.**
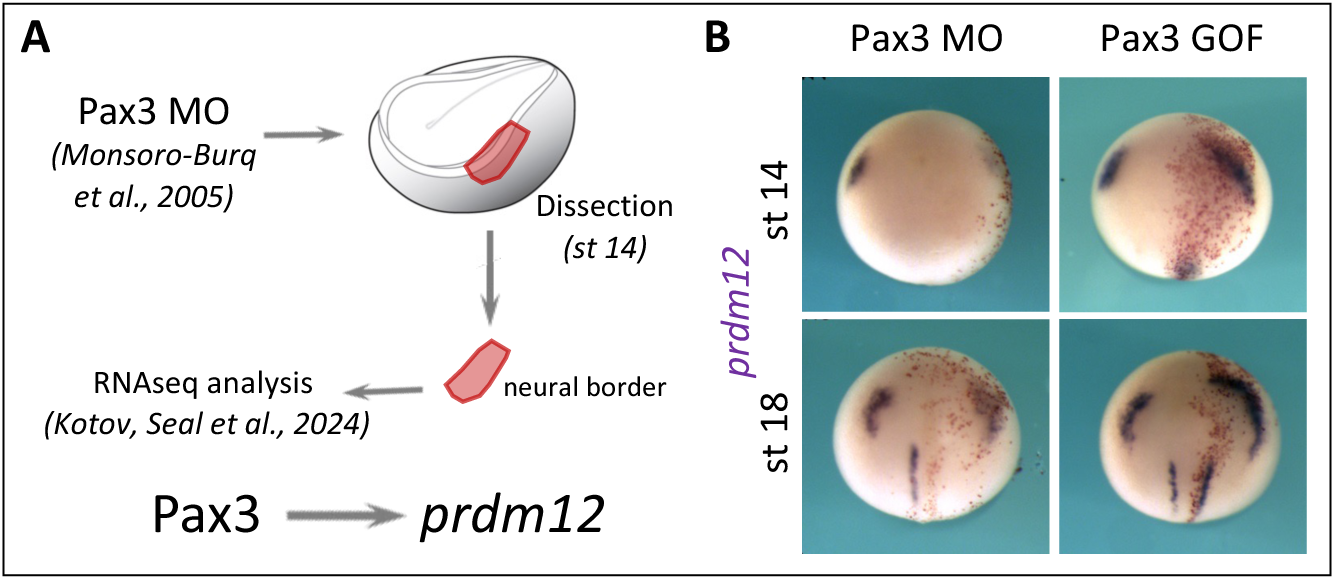
Prdm12 is a transcriptional target of Pax3. **(A)** Prdm12 was identified as a transcriptional target of the NB gene *pax3*. **(B)** Depletion or gain-of-function of *pax3* leads to a concomitant decrease or increase in the expression levels of *prdm12*, respectively. Red staining depicts the injected side. For st 14, effect = 26/26 (MO), 27/41 (GOF). For st 18, effect = 48/48 (MO), 18/51 (GOF).

**Fig. S2.**
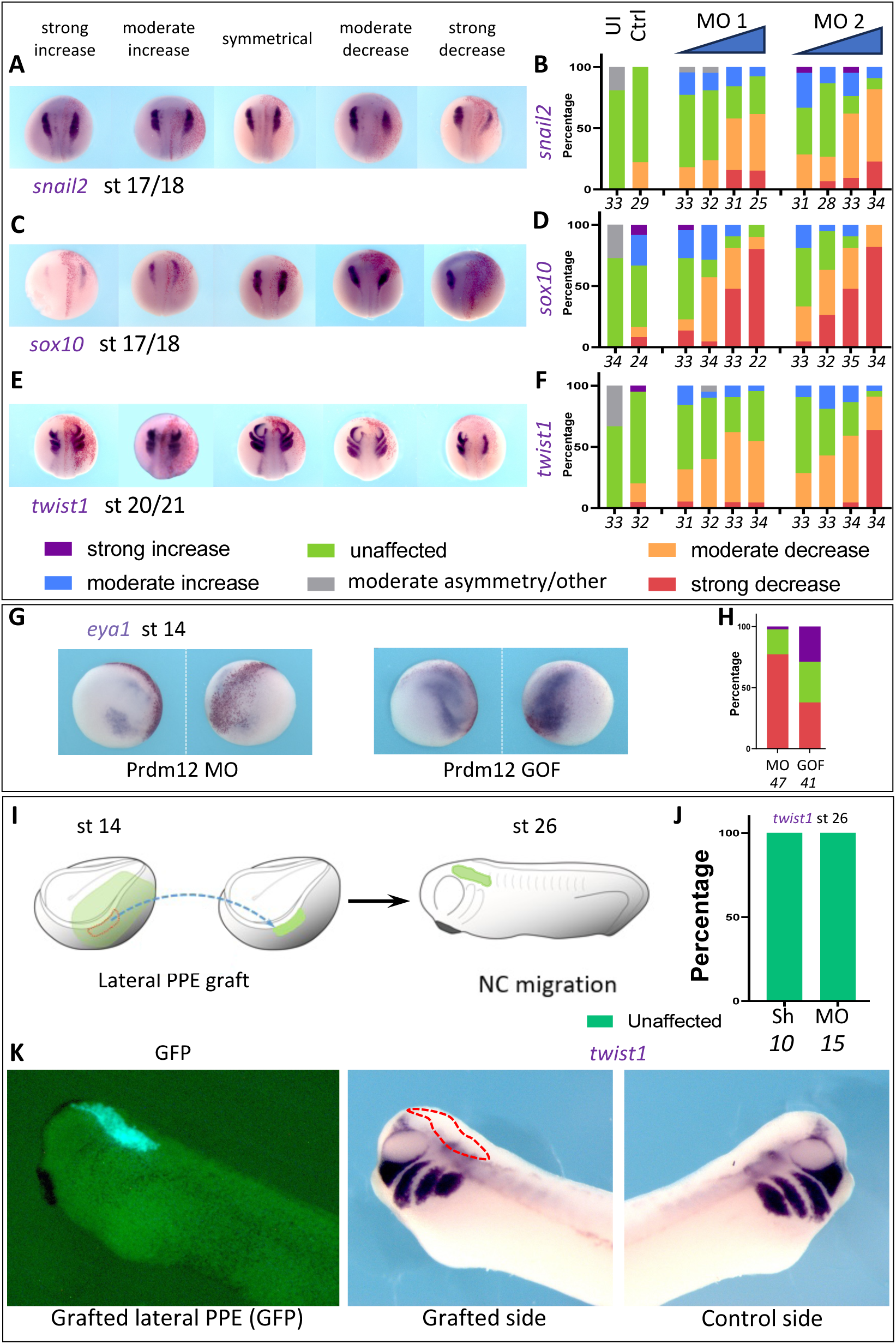
Prdm12 is required for NC-EMT and migration cell-autonomously. **(A, C, E)** ISH shows the range of phenotypes observed upon *prdm12* depletion upon *snail2* (A), *sox10* (C) and *twist1* (E). **(B, D, F)** Quantification of phenotypes shown in A, C and E respectively. UI – uninjected, Ctrl – control, MO1 – *prdm12* MO from *Matsukawa et al., 2015*, MO2 – *prdm12* MO from *Thélie et al., 2015*. **(G)** ISH shows the effect of *prdm12* upon *eya1* expression in the preplacodal ectoderm. **(H)** Quantification of phenotypes shown in (G). Legend as in (A-F). **(I)** Scheme of orthotopic lateral preplacodal ectoderm (PPE) graft. **(J, K)** NC migration phenotypes were observed by ISH against *twist1*. Quantification (J) of phenotypes observed (K) after lateral PPE graft. Sh – control lateral PPE graft, MO – *prdm12* morphant lateral PPE graft. Red dotted shape shows the graft location. In A, C, E and G, red staining depicts the injected side. Numbers under graphs depict number of embryos for each condition.

**Fig. S3.**
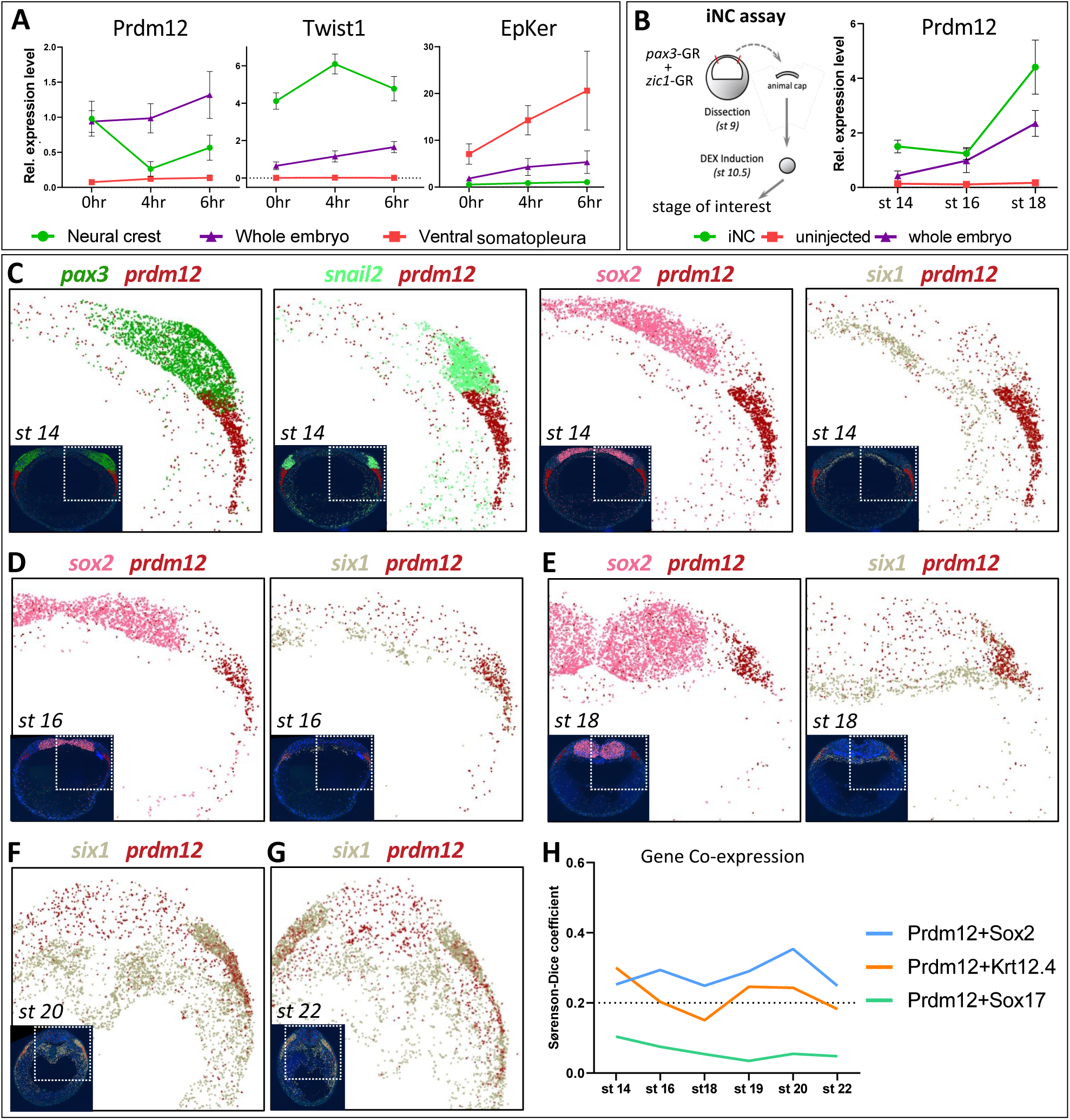
Expression of Prdm12 in the NC. **(A)** RT-qPCR experiment depicts the normalized relative expression of *prdm12*, *twist1* and epidermal keratin (*krt12.4*) in NC explants, the ventral somatopleura (negative control), and the whole embryo. Explants were microdissected at stage 17/18 and collected at different time points (grown at 18°C). N = 4. **(B)** Scheme representing the iNC assay. **(C)** RT-qPCR experiment depicts the normalized relative expression of *prdm12* in iNC explants collected at different stages. N = 6-7. **(C-G)** Spatial transcriptomics experiment depicts the expression patterns of different genes in transverse sections of embryos at different stages – (C) stage 14, (D) stage 16, (E) stage 18, (F) stage 20 and (G) stage 22. Main images depict the right dorsal side, inset images depict the whole section. **(H)** Sørenson-Dice coefficient depicts the degree of coexpression of different genes: x<0.2 – non-significant, 0.2<x<0.5 – moderate coexpression, 0.5<x – strong coexpression. (H) *Prdm12* shows low levels of overlap with *sox2* (neural ectoderm) and *krt12.4* (non-neural ectoderm), but not *sox17* (endoderm).

**Fig. S4.**
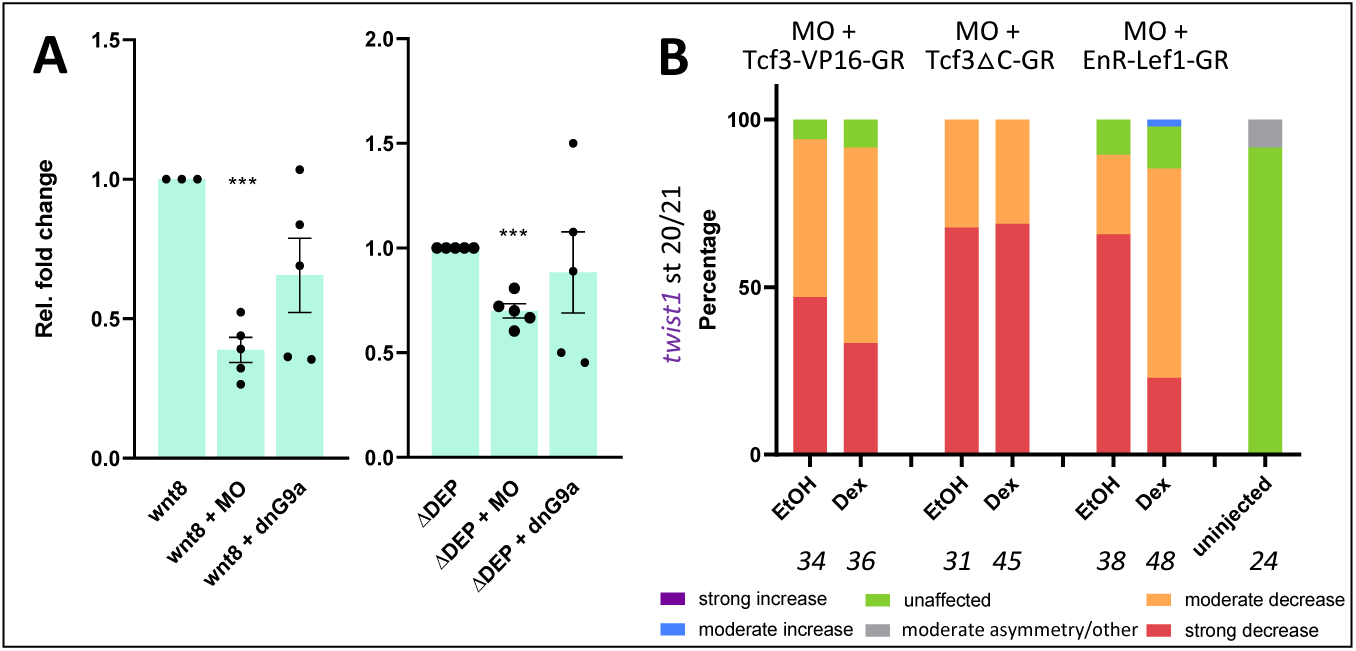
Effect of Prdm12 depletion on canonical Wnt signaling. **(A)** TOPFLASH-Luciferase assay shows the effect of *prdm12* depletion on canonical Wnt signaling readouts in animal cap explants microdissected stage 9 and grown till stage 17/18. MO – *prdm12* knockdown, ΔDEP – Dvl2 without the DEP domain, dnG9a – dominant-negative G9a. N ≥ 3. **(B)** ISH experiment against *twist1* at stage 20/21 depicts the effects of canonical Wnt (cWnt) modulation on NC migration in *prdm12* morphant embryos. GR-constructs used here can be translocated to the nucleus by treatment with Dexamethasone. Tcf3-VP16-GR – activation of cWnt, Tcf3△C-GR and EnR-Lef1-GR – inhibition of cWnt. Numbers under the graph depict number of embryos for each condition.

**Table S1.** Experimental reagents.

| Reagent | Experiment | Stage | Use | Amount | Reference |
| --- | --- | --- | --- | --- | --- |
| **Pax3 MO** | ISH | 2-cell | 1 cell | 20 ng | Monsoro-Burq et al., 2005 |
| **Prdm12 MO1** | ISH | 4-cell | 1 cell | 10/20/30/40 ng | Matsukawa et al., 2015 |
| **Prdm12 MO2** | ISH | 4-cell | 1 cell | 10/20/30/40 ng | Thélie et al., 2015 |
|  | iNC | 2-cell | 2 cells | 40 ng |  |
|  |  | 4-cell | 4 cells | 20 ng |  |
|  | Imaging, IF | 4-cell | 1 cell | 20 ng |  |
|  | Rescue (ISH, IF) | 8-cell | 1 cell | 20 ng |  |
| **Control MO** | ISH | 4-cell | 1 cell | 20 ng | Gene Tools, LLC |
|  | Imaging, IF | 4-cell | 1 cell | 20 ng |  |
| **Pax3-GR** | iNC | 2-cell | 2 cells | 30 pg | Milet et al., 2013 |
| **Zic1-GR** | iNC | 2-cell | 2 cells | 70 pg | Milet et al., 2013 |
| **Prdm12** | ISH | 4-cell | 1 cell | 500 pg | Thélie et al., 2015 |
|  | iNC | 2-cell | 2 cells | 1 ng |  |
| **dnG9a** | ISH | 4-cell | 1 cell | 500 pg | Thélie et al., 2015 |
|  | AC (Wnt) | 2-cell | 2 cells | 250 pg |  |
| **LA-mChe** | Imaging | 4-cell | 1 cell | 25 pg | Arnold et al., 2019 |
| **sf9-mNG** | Imaging | 4-cell | 1 cell | 25 pg | Arnold et al., 2019 |
| **CAAX-miRFP670** | Imaging | 4-cell | 1 cell | 25 pg | Arnold et al., 2019 |
| **ch N-Cadh** | Rescue (ISH) | 8-cell | 1 cell | 500 pg | Scarpa et al., 2015 |
| **dvl2 (all constructs)** | Rescue (ISH, IF) | 8-cell | 1 cell | 150 pg | Kratzer et al., 2020 |
|  | AC (Wnt) | 2-cell | 2 cells | 300 pg |  |
| **TOPFLASH-Luc** | AC (Wnt) | 2-cell | 2 cells | 25 pg | Promega |
| **Atf2-Luc** | AC (Wnt) | 2-cell | 2 cells | 25 pg | Ohkawara and Niehrs, 2011 |
| **Renilla-Luc** | AC (Wnt) | 2-cell | 2 cells | 5 pg | Promega |
| **wnt5** | AC (Wnt) | 2-cell | 2 cells | 30 pg | Monsoro-Burq et al., 2005 |
| **wnt8** | AC (Wnt) | 2-cell | 2 cells | 30 pg | Monsoro-Burq et al., 2005 |
| **tcf3-VP16-GR** | Rescue (ISH) | 8-cell | 1 cell | 150 pg | Maj et al., 2016 |
| **tcf3ΔC-GR** | Rescue (ISH) | 8-cell | 1 cell | 150 pg | Maj et al., 2016 |
| **enR-Lef1-GR** | Rescue (ISH) | 8-cell | 1 cell | 150 pg | Maj et al., 2016 |
| **GFP** | Imaging, IF | 4-cell | 1 cell | 250 pg | Milet and Monsoro-Burq, 2014 |
|  | iNC, AC (Wnt) | 2-cell | 2 cells | 500 pg |  |
| **β-gal** | ISH | 2-cell | 1 cell | 500 pg | Monsoro-Burq et al., 2005 |
|  |  | 4-cell | 1 cell | 250 pg |  |
|  |  | 8-cell | 1 cell | 250 pg |  |

**Table S2.**
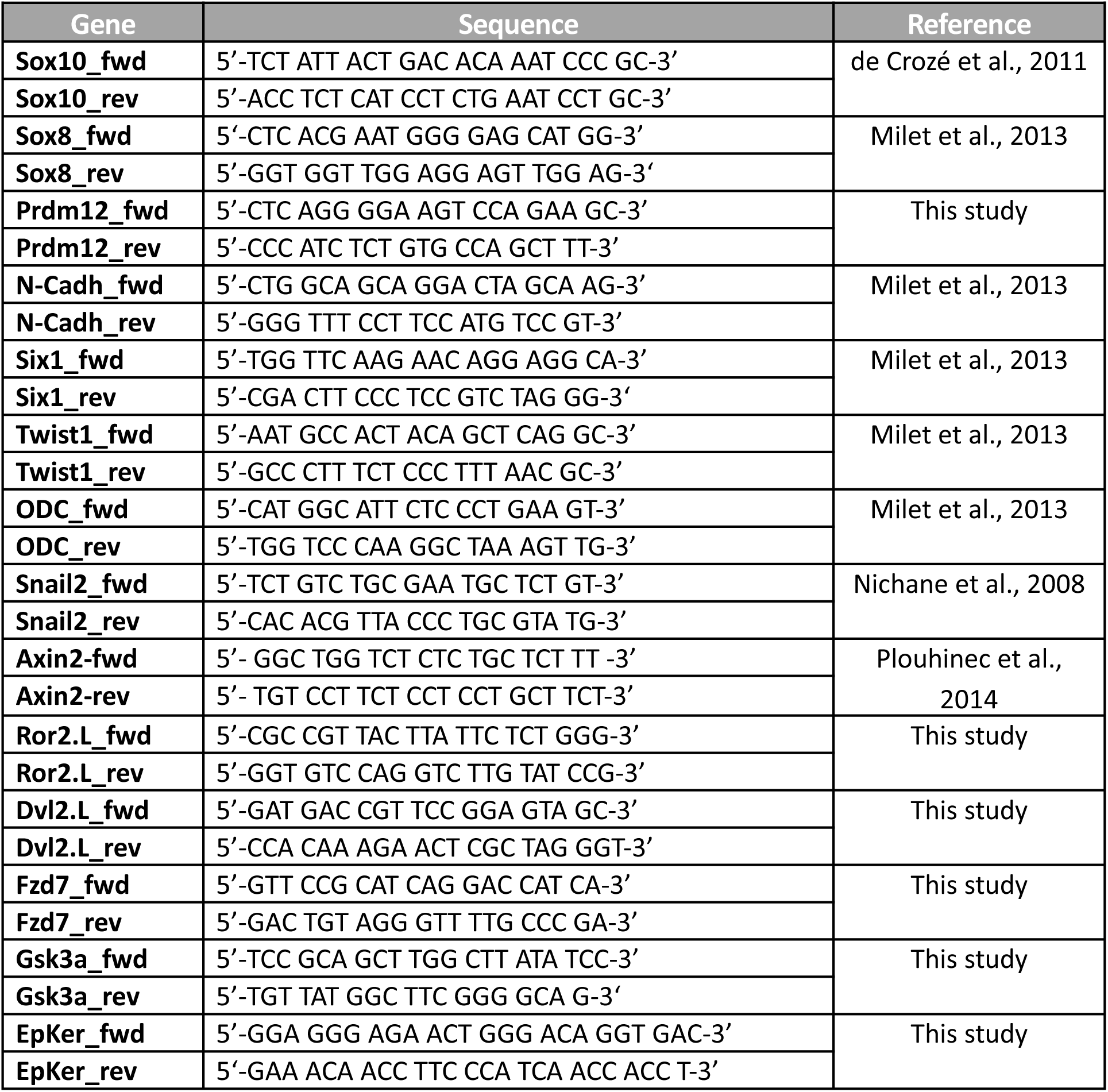
Primers used in this study for RT-qPCR.

